# Isolation of gametes and zygotes from *Setaria viridis*

**DOI:** 10.1101/2021.11.01.466850

**Authors:** Erika Toda, Takatoshi Kiba, Norio Kato, Takashi Okamoto

## Abstract

*Setaria viridis*, the wild ancestor of foxtail millet (*Setaria italica*), is an effective model plant for larger C_4_ crops because it has several desirable traits, such as short generation time, prolific seed production and a small genome size. These advantages are well suited for investigating molecular mechanisms in angiosperms, especially C_4_ crop species. Here, we report a procedure for isolating gametes and zygotes from *S. viridis* flowers. To isolate egg cells, ovaries were harvested from unpollinated mature flowers and cut transversely, which allowed direct access to the embryo sac. Thereafter, an egg cell was released from the cut end of the basal portion of the dissected ovary. To isolate sperm cells, pollen grains released from anthers were immersed in a mannitol solution, resulting in pollen-grain bursting, which released sperm cells. Additionally, *S. viridis* zygotes were successfully isolated from freshly pollinated flowers. Isolated zygotes cultured in a liquid medium developed into globular-like embryos and cell masses. Thus, isolated *S. viridis* gametes, zygotes and embryos are attainable for detailed observations and investigations of fertilization and developmental events in angiosperms.

## Introduction

In angiosperms, fertilization and subsequent developmental events, such as zygotic development/embryogenesis and endosperm development, occur in embryo sacs that are deeply embedded in ovular tissues (Nawaschin, 1898; Guignard, 1899; Russell, 1992; Raghavan, 2003). Consequently, investigations of fertilization and/or post-fertilization events in embryo sacs have been impeded by the inability to easily and directly analyze embedded female gametophytes. To overcome these difficulties, isolated gametes or zygotes have been used for analyzing the reproductive and developmental processes in embryo sacs. In the late 1980s, technical advances led to the successful isolation of viable gametes of angiosperm species (reviewed in Theunis *et al.*, 1991). In addition to gamete isolation, fertilized egg cells (zygotes) have been successfully isolated in cereal crops, such as barley (Holm *et al.*, 1994), maize (Leduc *et al.*, 1996), wheat (Kumlehn *et al.*, 1997, 1998) and rice (Zhang *et al.*, 1999). Furthermore, over the last 15 years, isolated gametes, zygotes or embryos were successfully used for transcriptome analyses to identify genes expressed specifically or preferentially in female gametes, male gametes, zygotes and early embryos (Sprunck *et al.*, 2005; Ning *et al.*, 2006; Yang *et al.*, 2006; Steffen *et al.*, 2007; Borges *et al.*, 2008; Wang *et al.*, 2010; Wuest *et al.*, 2010; Ohnishi *et al.*, 2011; Abiko *et al.*, 2013a; Anderson *et al.*, 2013; Anderson *et al.*, 2017; Chen *et al.*, 2017; Rahman *et al.*, 2019; Zhao *et al.*, 2019; Zhao *et al.*, 2020). The identified genes may be involved in reproductive or developmental processes, such as gamete differentiation, gamete fusion and early zygotic development. In addition to transcriptome analyses, proteins expressed in rice gametes were identified using proteomic analyses (Abiko *et al.*, 2013b).

Green foxtail millet (*Setaria viridis*), which is the wild ancestor of foxtail millet (*Setaria italica*), is an effective model for larger C_4_ panicoid crops, such as switchgrass, guinea grass, maize and sorghum (reviewed in Li and Brutnell, 2011). Because *S. viridis* has several advantageous traits, such as a short generation time, simple growth requirements, prolific seed production and a diploid with a relatively small genome size of approximately 510 Mb (Brutnell *et al.*, 2010; Li and Brutnell, 2011), it has recently been recognized as a new monocotyledonous model species that is now available in many laboratories. Therefore, gametes, zygotes and embryos isolated from *S. viridis* plants represent suitable materials for investigating the cellular and molecular mechanisms involved in zygote development and subsequent embryogenesis. Here, we established procedures in *S. viridis* for the isolation of gametes from unpollinated flowers and zygotes from pollinated flowers. In addition, the isolated *S. viridis* zygotes were cultured and their developmental profiles were observed.

## Materials and methods

### Plant materials

Seeds of *Setaria viridis* (accession A10.1) were provided by the Brutnell Laboratory (Brutnell *et al.*, 2010). The mature seeds were planted in soil and grown in an environmental chamber (Nippon Medical and Chemical Instruments) at 26°C/24°C with a 13-h light/11-h dark photoperiod.

### Isolation of gametes

For egg cell isolation, ovaries were harvested from unpollinated mature flowers using forceps and transferred to plastic dishes (φ 35 mm) containing 3 ml mannitol solution (370 mOsmol kg^−1^ H_2_O). After the ovaries were cut transversely using a thin razor blade, egg cells were released from the cut end of the basal portions of the dissected ovaries. To isolate sperm cells, anthers were harvested from unpollinated mature flowers and transferred to plastic dishes containing 3 ml mannitol solution (370 mOsmol kg^−1^ H_2_O). The anthers were broken in the mannitol solution using forceps to release the pollen grains, and thereafter, the pollen grains burst and released their contents, including sperm cells.

### Isolation of zygotes

For zygote isolation, ovaries were harvested from pollinated flowers having fresh stigmata, which indicated that they had been pollinated on the day of harvest, and transferred to plastic dishes containing 3 ml mannitol solution (370 mOsmol kg^−1^ H_2_O). To collect zygotes efficiently, enlarged ovaries, in which embryogenesis and endosperm development had already progressed, were removed during the ovary-harvesting process. The isolated ovaries were cut as described above, and the cut ovaries were incubated in mannitol solution for approximately 1–2 h. Thereafter, the zygotes were released by gently pushing the cut end of the basal portions of dissected ovaries with a glass needle.

### Culturing zygotes

Isolated zygotes were transferred into a 1-μl droplet of mannitol solution (370 mOsmol kg^−1^ H_2_O) overlaid with mineral oil on a coverslip (Okamoto, 2011) and washed three times by gently transferring the cells into droplets of fresh mannitol solution (450 mOsmol kg^−1^ H_2_O). Thereafter, the isolated zygotes were cultured in a Millicell-CM insert (Merck Millipore) containing N6Z (Uchiumi *et al.*, 2007) or MSO (Kranz and Lörz, 1993) medium as described by Okamoto (2011). Zygotes were cultured at 26°C in darkness.

## Results and Discussion

### Developmental stages of *S. viridis* flowers

We first harvested panicles from *S. viridis* plants (Fig. 1a and 1b). Thereafter, *S. viridis* flowers at various developmental stages were harvested and dissected to determine the stages suitable for the isolation of gametes and zygotes. Figure 1c shows an unpollinated *S. viridis* flower. When the unpollinated flowers were dissected, immature ovaries with undeveloped anthers and stigmata (Fig. 1d and 1e) or putative mature ovaries with developed anthers and stigmata (Fig. 1f) were isolated. Pollination occurs when flowers open, and the flowers close again after pollination. Thereafter, the stigma remaining on the flower (Fig. 1g) is an indication of a pollinated flower. The pollinated flowers with fresh stigmata were harvested (Fig. 1g), and the ovaries were isolated from the pollinated flowers (Fig. 1h). The ovaries from the pollinated flowers having fresh stigmata were similar in size to unpollinated mature ovaries (Fig. 1f). Additionally, pollinated flowers with degenerated stigmata were harvested (Fig. 1i), and developing ovaries were isolated from the pollinated flowers (Fig. 1j and 1k). The flowering of *S. viridis* spikelets is triggered by the darkness of night and low temperatures (Rizal *et al.*, 2013). Therefore, it was presumed that the flowers having fresh stigmata (Fig. 1g) were pollinated on the day of harvest, and that these freshly pollinated flowers would be suitable for isolating zygotes.

**Figure 1.**
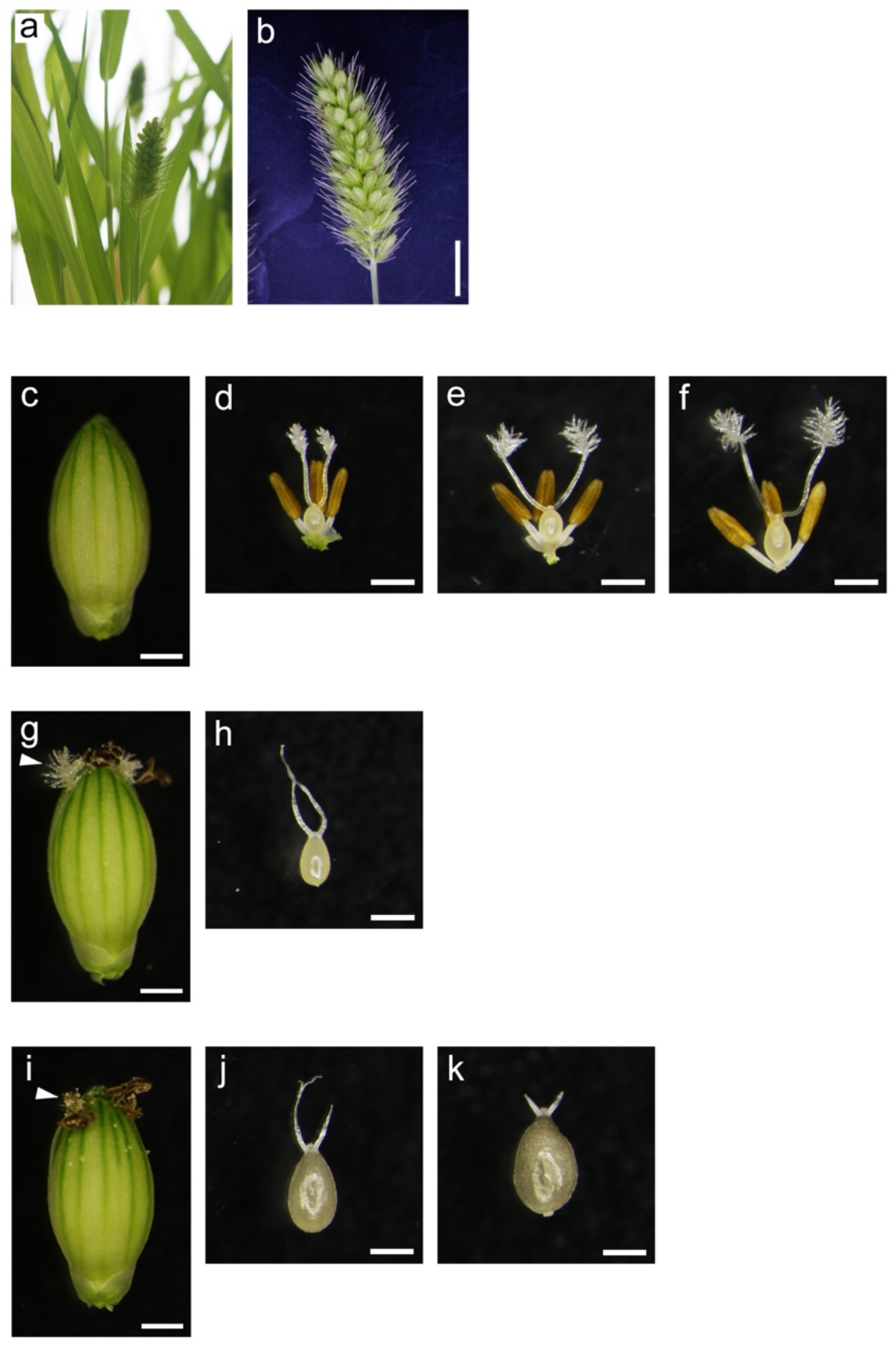
Female and male reproductive organs harvested from unpollinated and pollinated *S. viridis* flowers. (a) *S. viridis* plants. (b) A panicle. (c) An unpollinated flower. (d–f) Immature (d and e) and mature (f) pistils and anthers harvested from unpollinated flowers. (g) A pollinated flower having a fresh stigma. (h) An ovary harvested from a pollinated flower having a fresh stigma. (i) A pollinated flower having a degraded stigma. (j and k) Ovaries harvested from the pollinated flowers. Stigmata and anthers of pollinated flowers were removed during the ovary isolation process. Arrowheads in panels (g) and (i) indicate fresh and degenerated stigmata, respectively. Bars = 5 mm in (b) and 0.5 mm in (c–k).

### Isolation of *S. viridis* egg cells

For egg cell isolation, mature ovaries harvested from the unpollinated flowers (Fig. 2a) were transferred into a mannitol solution adjusted to 370 mOsmol kg^−1^ H_2_O, as in the procedure for isolating rice egg cells (Uchiumi *et al.*, 2006), and the ovaries were cut transversely in the mannitol solution (Fig. 2b). Putative egg cells were released from the cut end of the basal portions of the dissected ovaries (Fig. 2c and 2d). Furthermore, an egg apparatus, consisting of an egg cell and two synergid cells, which is a characteristic feature of angiosperms, was also released (Fig. 2e). The *S. viridis* egg cells and synergid cells were approximately 30 μm in diameter (Fig. 2e), and the isolated putative egg cells (Fig. 2c and 2d) were almost similar in size to the egg cells in the egg apparatuses (Fig. 2e). The *S. viridis* egg cells were smaller than those of maize (60–77 μm; Kranz *et al.*, 1991), wheat (50–70 μm; Kovacs *et al.*, 1994) and rice (40–50 μm; Uchiumi *et al.*, 2006), but were similar to those of Brachypodium (30–35 μm; Matsumura and Okamoto, 2016). In addition, the isolated *S. viridis* egg cells showed many putative vacuoles, ranging in size from approximately 6 to 10 μm, that existed at their peripheral regions (Fig. 2e). The peripheral localization of vacuoles is consistent with the cellular characteristics of egg cells isolated from maize (Faure *et al.*, 1992), wheat (Kovacs *et al.*, 1994), rice (Uchiumi *et al.*, 2007) and Brachypodium (Matsumura and Okamoto, 2016), suggesting that the cells isolated from the ovaries are egg cells.

**Figure 2.**
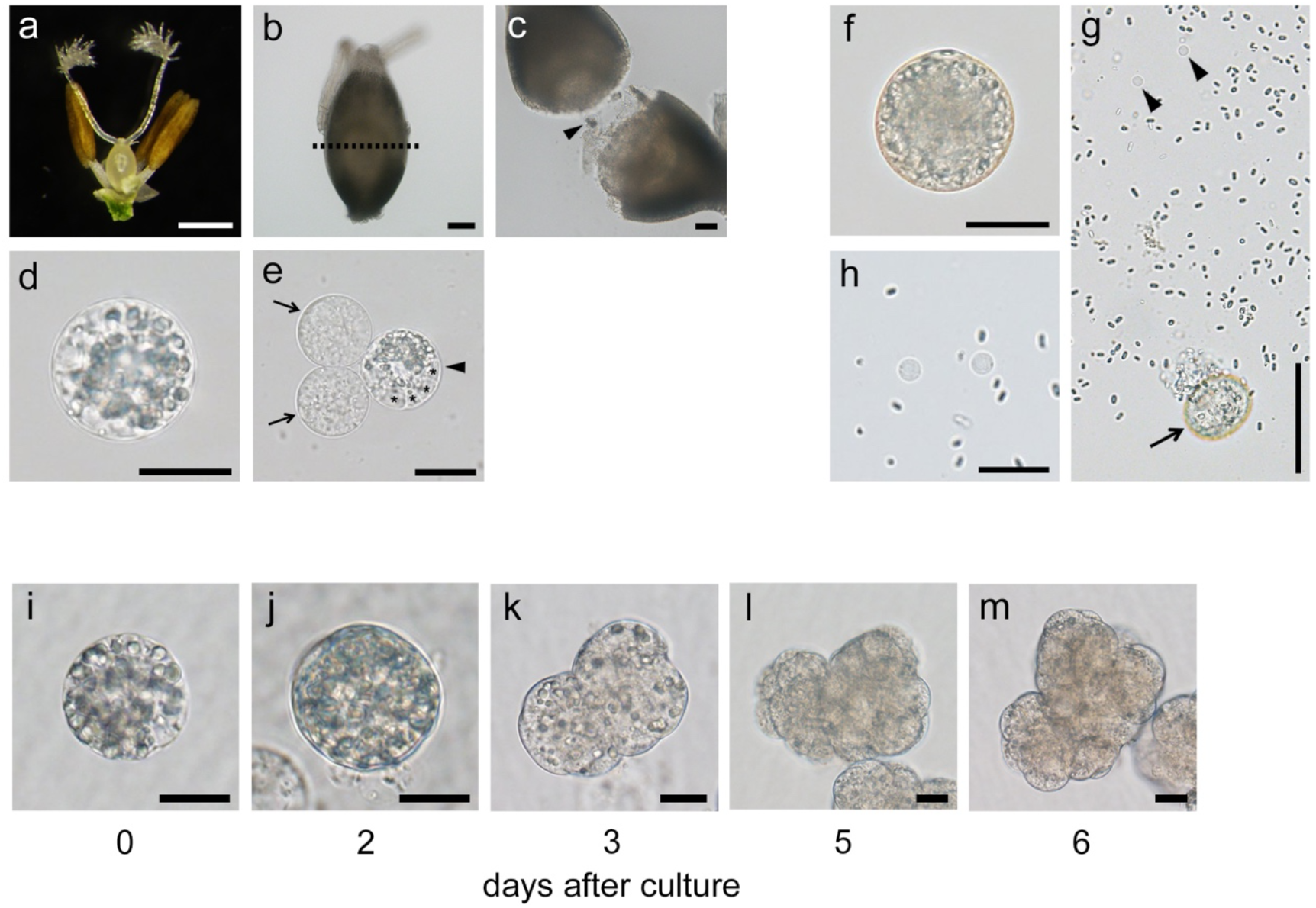
Isolation of *S. viridis* gametes and the developmental profiles of zygotes isolated from pollinated flowers. (a) An ovary harvested from unpollinated mature flower. (b) An isolated ovary. (c) An egg cell (arrowhead) released from the cut end of the basal portion of a dissected ovary. (d) An isolated egg cell. (e) An egg apparatus consisting of an egg cell and two synergid cells. (f) A pollen grain. (g) Two sperm cells released from a pollen grain. (h) Isolated sperm cells. After 2 days of culturing, the isolated zygote (i) developed into a globular-like embryo (j) and after 3, 5 and 6 days of culturing, a growing cell mass was observed (k–m). The dotted black line in (b) indicates the cutting line on the ovary for egg cell isolation. Arrowheads in (c) and (e) indicate the isolated egg cells and the egg cell of an egg apparatus, respectively. Arrows in (e) indicate the synergid cells of an egg apparatus. Asterisks in (e) indicate putative vacuoles in the egg cell. The arrow and arrowheads in (g) indicate a pollen grain releasing its contents and the released sperm cells, respectively. Bars = 0.5 mm in (a), 100 μm in (b), 50 μm in (c) and (g), 20 μm in (d–f), (h) and (i–m).

### Isolation of *S. viridis* sperm cells

When pollen grains are soaked in an appropriate osmotic solution, they generally burst and release their contents, including sperm cells (Kranz *et al.*, 1991; Theunis *et al.*, 1991). Therefore, we applied the osmotic bursting technique to isolate sperm cells from *S. viridis* pollen grains. Anthers were harvested from unpollinated mature flowers (Fig. 2a) and transferred into a mannitol solution adjusted to 370 mOsmol kg^−1^ H_2_O. The anthers were broken in the mannitol solution using forceps to release the pollen grains (Fig. 2f). After 5–10 min of the pollen grains being immersed in the mannitol solution, the grains burst and released their contents, including two sperm cells (Fig. 2g) because *S. viridis* plants form tricellular-type pollen grains. The *S. viridis* pollen grains and sperm cells were approximately 40 μm and 5 μm in diameter, respectively (Fig. 2f and 2h). Notably, the pollen grains isolated from immature unpollinated flowers (Fig. 1d and 1e) rarely burst and released their contents (data not shown), indicating that mature pollen grains are appropriate for the isolation of *S. viridis* sperm cells. Previously, *S. viridis* sperm cells had been successfully isolated through the immersion of pollen grains in a mannitol solution and used for *in vitro* fusions with isolated wheat gametes (Li *et al.*, 2019). In Li *et al.* (2019), the sperm cells were released in 6%–12% mannitol solutions. We also successfully isolated *S. viridis* sperm cells using a mannitol solution adjusted to 370 mOsmol kg^−1^ H_2_O (approximately 6% concentration), suggesting that *S. viridis* sperm cells adapt to a wide osmolarity range.

### Isolation of *S. viridis* zygotes and their developmental profiles

The zygotes (1-celled embryos) were isolated from pollinated flowers having fresh stigmata (Fig. 1g). The isolated ovaries (Fig. 1h) were transferred into a mannitol solution (370 mOsmol kg^−1^ H_2_O) and cut transversely using a thin razor blade, as in the isolation of *S. viridis* egg cells. Thereafter, the dissected ovaries were incubated in the mannitol solution for approximately 1–2 h because the putative zygotes easily burst just after the ovaries were dissected. After the incubation, a putative zygote was released from the cut end of the basal portion of each dissected ovary by gently pushing with a glass needle (Fig. 2i). To monitor their developmental profiles, the isolated putative *S. viridis* zygotes were cultured in a Millicell-CM insert containing the N6Z culture medium, which has been used for culturing zygotes of barley (Kumlehn *et al.*, 1999), rice (Uchiumi *et al.*, 2007) and wheat (Maryenti *et al.*, 2019). After 2 days of culturing, the cells developed into globular-like embryos (Fig. 2j) and continued undergoing cell division, forming cell masses, until approximately 6 days after culturing began (Fig. 2k–m). This indicated that the cells isolated from pollinated flowers are zygotes capable of developing into embryos. The developmental ratio of the isolated zygotes was 35.7% (n = 10/28; Table 1).

**Table 1.**
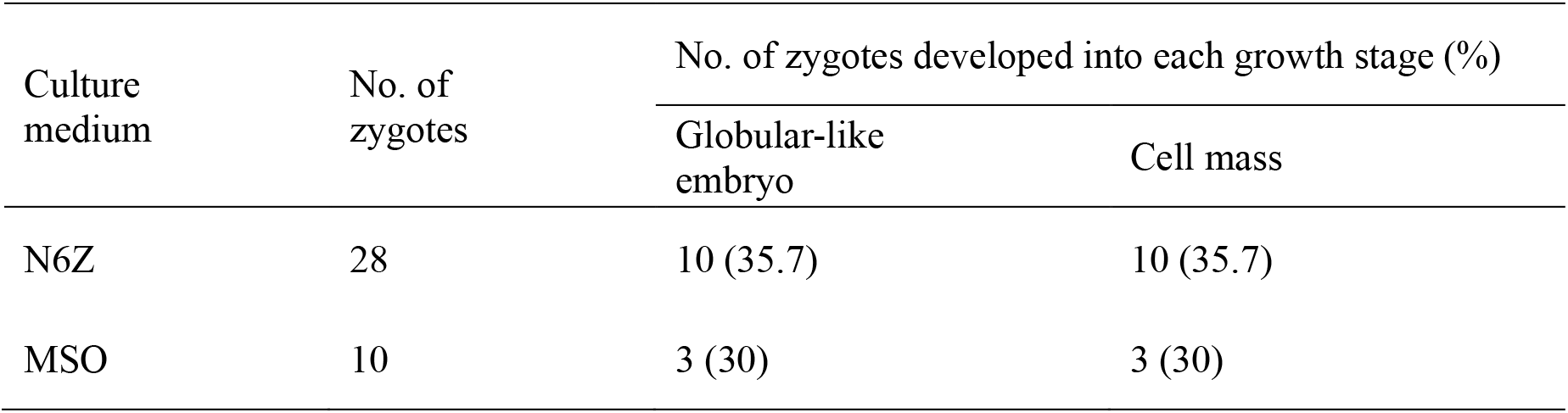
Developmental profiles of isolated *S. viridis* zygotes

Although the zygotes divided and reached the cell mass stage, the development of the cell masses was arrested. Therefore, we cultured the isolated *S. viridis* zygotes in MSO medium, which has been used for culturing maize zygotes (Kranz and Lörz, 1993) because *S. viridis*, like maize, is C_4_ crop species. However, the ratio of the isolated zygotes that developed to the cell mass stage in the MSO medium was similar to that in the N6Z medium (n = 3/10, Table 1), and the cell masses also underwent developmental arrest. Because the shape of the isolated putative zygotes (Fig. 2i) was similar to that of the isolated egg cells (Fig. 2d), it was difficult to discriminate precisely between the two and to isolate only zygotes in which fertilization had successfully occurred. Therefore, the developmental ratio may have been affected by the contamination of egg cells isolated from pollinated flowers with unsuccessful gamete fusions.

### Uses of isolated gametes and zygotes in further investigations

In the present study, we established procedures in *S. viridis* for the isolation of gametes from unpollinated mature flowers and of zygotes from pollinated flowers. Moreover, the isolated zygotes were cultured in a liquid medium, resulting in their successful development into globular-like embryos and cell masses. To observe and analyze the detailed processes of fertilization and post-fertilization in angiosperms, such as karyogamy (Faure *et al.*, 1993; Ohnishi *et al.*, 2014), male chromatin decondensation in zygotes (Scholten *et al.*, 2002), zygotic initiation (Rahman *et al.*, 2019) and development (Kranz *et al.*, 1995), parental genome contributions to zygotic development (Toda *et al.*, 2016, 2018) and fertilization-induced or - suppressed gene expression (Okamoto *et al.*, 2005; Rahman *et al.*, 2019), isolated gametes have been used to produce a zygote through the *in vitro* fusion of an egg cell and a sperm cell. Using isolated gametes, complete *in vitro* fertilization (IVF) systems have been established in three major crop species, maize (Kranz and Lörz, 1993), rice (Uchiumi *et al.*, 2007) and wheat (Maryenti *et al.*, 2019). An IVF system using the present gamete-isolating techniques would allow the investigation of zygotic development and embryogenesis directly in *S. viridis*, although further culture condition improvements will be required to obtain plantlets from isolated zygotes. Furthermore, the isolated gametes and zygotes may be used for a wide range of single cell-type omics analyses. Thus, the present procedures for isolating *S. viridis* gametes and zygotes are widely applicable to the investigation of cellular and molecular mechanisms involved in fertilization processes, post-fertilization events, zygotic development and embryogenesis in angiosperms, particularly in C_4_ crop species.

## Acknowledgements

We thank Dr. T. P. Brutnell (Chinese Academy of Agricultural Sciences, China) for providing seeds of *S. viridis*, Ms. A. Takebayashi (RIKEN, Japan) for preparing *S. viridis* plants, and the RIKEN Bio Resource Center (Tsukuba, Japan) for providing cultured rice cells (Oc line).

## Ethical standards

Not applicable.

## References

Abiko M, Furuta K, Yamauchi Y, Fujita C, Taoka M, Isobe T, Okamoto T (2013b) Identification of proteins enriched in rice egg or sperm cells by single-cell proteomics. PLoS ONE 8, e69578.

Abiko M, Maeda H, Tamura K, Hara-Nishimura I, Okamoto T (2013a) Gene expression profiles in rice gametes and zygotes: Identification of gamete-enriched genes and up- or down-regulated genes in zygotes after fertilization. J Exp Bot 64, 1927–1940.

Anderson SN, Johnson CS, Jones DS, Conrad LJ, Gou X, Russell SD, Sundaresan V (2013) Transcriptomes of isolated *Oryza sativa* gametes characterized by deep sequencing: Evidence for distinct sex-dependent chromatin and epigenetic states before fertilization. Plant J 76, 729–741.

Anderson SN, Johnson CS, Chesnut J, Jones DS, Khanday I, Woodhouse M, Li C, Conrad LJ, Russell SD, Sundaresan V (2017) The zygotic transition is initiated in unicellular plant zygotes with asymmetric activation of parental genomes. Dev Cell 43, 349–358.

Borges F, Gomes G, Gardner R, Moreno N, McCormick S, Feijó JA, Becker JD (2008) Comparative transcriptomics of Arabidopsis sperm cells. Plant Physiol 148, 1168–1181.

Brutnell TP, Wang L, Swartwood K, Goldschmidt A, Jackson D, Zhu XG, Kellogg E, Eck JV (2010) *Setaria viridis*: a model for C_4_ photosynthesis. Plant cell 22, 2537–2544.

Chen J, Strieder N, Krohn NG, Cyprys P, Sprunck S, Engelmann JC, Dresselhaus T (2017) Zygotic genome activation occurs shortly after fertilization in maize. Plant Cell 29, 2106–2125.

Faure JE, Mogensen HL, Kranz E, Digonnet C, Dumas C (1992) Ultrastructural characterization and three-dimensional reconstruction of isolated maize (*Zea mays* L.) egg cell protoplasts. Protoplasma 171, 97–103.

Faure JE, Mogensen HL, Dumas C, Lörz H, Kranz E (1993) Karyogamy after electrofusion of single egg and sperm cell protoplasts from maize: cytological evidence and time course. Plant Cell 5, 747–755.

Guignard ML (1899) Sur les antherozoides et la double copulation sexuelle chez les vegetaux angiosperms. Rev Gén Bot 11, 129–135.

Holm PB, Knudsen S, Mouritzen P, Negri D, Olsen FL, Roue C (1994) Regeneration of fertile barley plants from mechanically isolated protoplasts of the fertilized egg cell. Plant Cell 6, 531–543.

Kovacs M, Barnabas B, Kranz E (1994) The isolation of viable egg cells of wheat (*Triticum aestivum* L.). Sex Plant Reprod 7, 311–312.

Kranz E, Bautor J, Lörz H (1991) *In vitro* fertilization of single, isolated gametes of maize mediated electrofusion. Sex Plant Reprod 4, 12–16.

Kranz E, Lörz H (1993) *In vitro* fertilization with isolated, single gametes results in zygotic embryogenesis and fertile maize plants. Plant Cell 5, 739–746.

Kranz E, von Wiegen P, Lörz H (1995) Early cytological events after induction of cell division in egg cells and zygote development following in vitro fertilization with angiosperm gametes. Plant J 8, 9–23.

Kumlehn J, Schieder O, Lörz H (1997) In vitro development of wheat (*Triticum aestivum* L.) from zygote to plant via ovule culture. Plant Cell Rep 16, 663–667.

Kumlehn J, Lörz H, Kranz E (1998) Differentiation of isolated wheat zygotes into embryos and normal plants. Planta 205, 327–333.

Kumlehn J, Lörz H, Kranz E (1999) Monitoring individual development of isolated wheat zygotes: a novel approach to study early embryogenesis. Protoplasma 208, 156–162.

Leduc N, Matthys-Rochon E, Rougier M, Mogensen L, Holm P, Magnard JL, Dumas C (1996) Isolated maize zygotes mimic *in vivo* embryonic development and express microinjected genes when cultured *in vitro*. Dev Biol 177, 190–203.

Li DX, Hu HY, Ru ZG, Tian HQ (2019) Wheat egg *in vitro* fusion with wheat and green bristlegrass sperm. Zygote 27, 126–130.

Li P, Brutnell TP (2011) *Setaria viridis* and *Setaria italica*, model genetic systems for the Panicoid grasses. J Exp Bot 62, 3031–3037.

Maryenti T, Kato N, Ichikawa M, Okamoto T (2019) Establishment of an *in vitro* fertilization system in wheat (*Triticum aestivum* L.). Plant Cell Physiol 60, 835–843.

Matsumura T, Okamoto T (2016) Isolation of gametes from *Brachypodium distachyon*. Plant Biotechnol 33, 39–43.

Nawaschin S (1898) Revision der Befruchtungsvorgange bei Lilium martagon und Fritillaria tenella. Bull Acad Imp Sci St Pétersbourg 9, 377–382.

Ning J, Peng X-B, Qu L-H, Xin HP, Yan TT, Sun MX (2006) Differential gene expression in egg cells and zygotes suggests that the transcriptome is restructed before the first zygotic division in tobacco. FEBS Lett 580, 1747–1752.

Ohnishi T, Takanashi H, Mogi M, Takahashi H, Kikuchi S, Yano K, Okamoto T, Fujita M, Kurata N, Tsutsumi N (2011) Distinct gene expression profiles in egg and synergid cells of rice as revealed by cell type-specific microarrays. Plant Physiol 155, 881–891.

Ohnishi Y, Hoshino R, Okamoto T (2014) Dynamics of male and female chromatin during karyogamy in rice zygotes. Plant Physiol 165, 1533–1543.

Okamoto T (2011) In vitro fertilization with rice gametes: production of zygotes and zygote and embryo culture. Methods Mol Biol 710, 17–27.

Okamoto T, Scholten S, Lörz H, Kranz E (2005) Identification of genes that are up- or down-regulated in the apical or basal cell of maize two-celled embryos and monitoring their expression during zygote development by a cell manipulation- and PCR-based approach. Plant Cell Physiol 46, 332–338.

Raghavan V (2003) Some reflections on double fertilization, from its discovery to the present. New Phytol 159, 565–583.

Rahman MH, Toda E, Kobayashi M, Kudo T, Koshimizu S, Takahara M, Iwami M, Watanabe Y, Sekimoto H, Yano K, Okamoto T (2019) Expression of genes from paternal alleles in rice zygotes and involvement of OsASGR-BBML1 in initiation of zygotic development. Plant Cell Physiol 60, 725–737.

Rizal G, Acebron K, Mogul R, Karki S, Larazo N, Quick WP (2013) Study of flowering pattern in *Setaria viridis*, a proposed model species for C_4_ photosynthesis research. J Bot 2013, 1–7.

Russell SD (1992) Double fertilization. Int Rev Cytol 40, 357–390.

Scholten S, Lörz H, Kranz E (2002) Paternal mRNA and protein synthesis coincides with male chromatin decondensation in maize zygotes. Plant J 32, 221–231.

Sprunck S, Baumann U, Edwards K, Langridge P, Dresselhaus T (2005) The transcript composition of egg cells changes significantly following fertilization in wheat (*Triticum aestivum* L.). Plant J 41, 660–672.

Steffen JG, Kang IH, Macfarlane J, Drews GN (2007) Identification of genes expressed in the *Arabidopsis* female gametophyte. Plant J 51, 281–292.

Theunis C, Pierson F, Cresti M (1991) Isolation of male and female gametes in higher plants. Sex Plant Reprod 4, 145–154.

Toda E, Ohnishi Y, Okamoto T (2016) Development of polyspermic rice zygotes. Plant Physiol 171, 206–214.

Toda E, Ohnishi Y, Okamoto T (2018) An imbalanced parental genome ratio affects the development of rice zygotes. J Exp Bot 69, 2609–2619.

Uchiumi T, Komatsu S, Koshiba T, Okamoto T (2006) Isolation of gametes and central cells from *Oryza sativa* L. Sex Plant Reprod 19, 37–45.

Uchiumi T, Uemura I, Okamoto T (2007) Establishment of an *in vitro* fertilization system in rice (*Oryza sativa* L.). Planta 226, 581–589.

Wang D, Zhang CQ, Hearn DJ, Kang IH, Punwani JA, Skaggs MI, Drews GN, Schumaker KS, Yadegari R (2010) Identification of transcription-factor genes expressed in the Arabidopsis female gametophyte. BMC Plant Biol 10, 110.

Wuest SE, Vijverberg K, Schmidt A, Weiss M, Gheyselinck J, Lohr M, Wellmer F, Rahnenführer J, von Mering C, Grossniklaus U (2010) Arabidopsis female gametophyte gene expression map reveals similarities between plant and animal gametes. Curr Biol 20, 506–512.

Yang H, Kaur N, Kiriakopolos S, McCormick S (2006) EST generation and analyses towards identifying female gametophyte-specific genes in *Zea mays* L. Planta 224, 1004–1014.

Zhang J, Dong WH, Galli A, Potrykus I (1999) Regeneration of fertile plants from isolated zygotes of rice (*Oryza sativa*). Plant Cell Rep 19, 128–132.

Zhao P, Zhou X, Shen K, Liu Z, Cheng T, Liu D, Cheng Y, Peng X, Sun MX (2019) Two-step maternal-to-zygotic transition with two-phase parental genome contributions. Dev Cell 49, 882–893.

Zhao P, Zhou X, Zheng Y, Ren Y, Sun MX (2020) Equal parental contribution to the transcriptome is not equal control of embryogenesis. Nat Plants 6, 1354–1364.

